# Caring without sharing: Meta-analysis 2.0 for massive genome-wide association studies

**DOI:** 10.1101/436766

**Authors:** Armin Pourshafeie, Carlos D. Bustamante, Snehit Prabhu

## Abstract

Genome-wide association studies have been effective at revealing the genetic architecture of simple traits. Extending this approach to more complex phenotypes has necessitated a massive increase in cohort size. To achieve sufficient power, participants are recruited across multiple collaborating institutions, leaving researchers with two choices: either collect all the raw data at a single institution or rely on meta-analyses to test for association. In this work, we present a third alternative. Here, we implement an entire GWAS workflow (quality control, population structure control, and association) in a fully decentralized setting. Our iterative approach (a) does not rely on consolidating the raw data at a single coordination center, and (b) does not hinge upon large sample size assumptions at each silo. As we show, our approach overcomes challenges faced by meta-studies when it comes to associating rare alleles and when case/control proportions are wildly imbalanced at each silo. We demonstrate the feasibility of our method in cohorts ranging in size from 2K (small) to 500K (large), and recruited across 2 to 10 collaborating institutions.

## Introduction

Genome wide association studies (GWAS) are a popular approach to elucidate genetic architecture of human phenotypes. This design has led to the discovery of many novel loci underpinning a panoply of human traits (see Visscher et. al. for a recent review (1)). For traits driven by few variants with large effects, moderately sized cohorts have been sufficient to power discovery. However, the GWAS framework demands increasingly larger cohort sizes as the complexity of the trait grows. To achieve required statistical power today, large, multi-institutional consortia are assembled under a common data sharing agreement. Meta-analysis (2; 3; 4) or a combination of meta- and mega-analysis (centralized analysis) (5; 6; 7; 8) constitute two major approaches to conducting GWAS.

Each approach offers merits and shortcomings. For mega-analysis, collecting all the data at every analysis core is not only expensive and time consuming but also creates a security vulnerability at ach institution that hosts a copy of the data. Conversely, the meta-analysis approach eliminates the need for data replication, but is more limited in flexibility. In particular, (a) subtle differences in models, assumptions, and quality control (QC) can introduce biases in the results (9), (b) the shared data and summary statistics might be inadequate for some types of inference (e.g. individual level population structure control, conditional or joint analysis, etc.), and (c) parameter estimates can be unreliable for rare variants or from centers contributing small sample sizes because the asymptotic properties of maximum likelihood estimation theory may not hold (10).

In this manuscript we develop a method that interpolates between centralized and meta-analysis methods. Like meta-studies, our paradigm leverages data fragmented across multiple silos, where each silo communicates aggregate level information with a central hub to perform the GWAS. Unlike meta-studies, in our paradigm the parameter estimates are computed iteratively (figure 1). To contrast these two paradigms, we henceforth refer to the one-time communication setting of traditional meta-analysis as “Meta 1.0”, and to our iterative setting as “Meta 2.0” (see Figure1).

**Figure 1:**
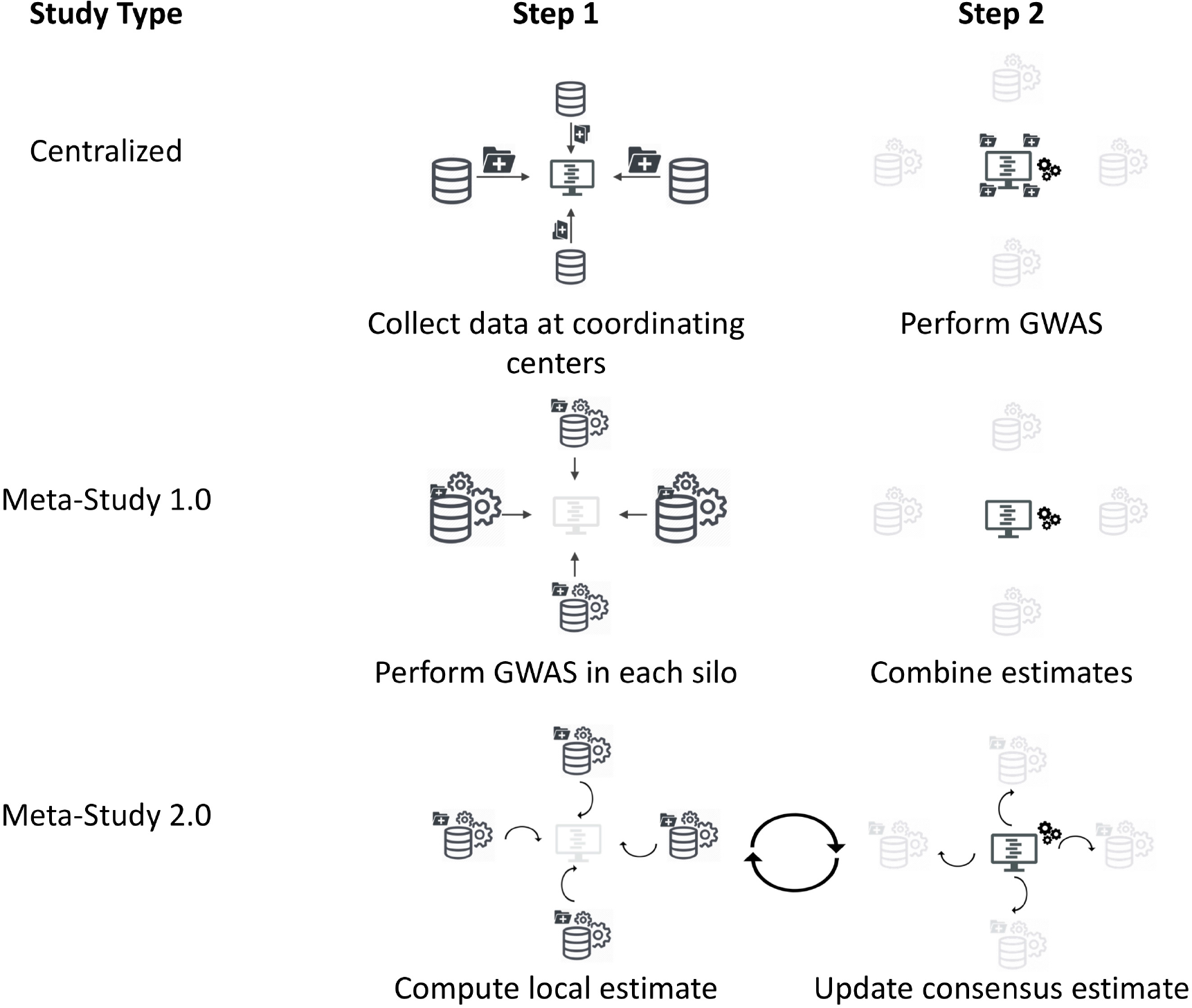
Comparison of data-flows. In the mega-study (centralized) framework, the data is gathered and analyzed at a single analysis center. For a typical meta-analysis approach (meta 1.0), summary statistics from previously analyzed datasets in collected and analyzed at the center. In our proposed framework (meta 2.0), the data is analyzed at each silo and a summary statistics is computed and reported to the center. Consequently, a global consensus is achieved at the center and reported back to each silo. The consensus estimates can then be used to improve the local estimates at each silo and the process can be iterated

We show that all steps of a canonical case-control GWAS can be implemented in our framework, including quality control (QC) filters and population structure control by regressing out the top principal components (PCs). In particular, typical meta-analysis assumptions on the distribution of participant attributes in each silo (e.g. case-control status, ancestry, etc.) can be relaxed while simultaneously attaining near-perfect parameter estimates. For GWAS, our pipeline leverages an algorithm for performing PCA by communicating the LD-matrices (Algorithm 1), and subsequently a method for performing generalized regression (most commonly, linear and logistic regression) by iteratively solving a regularized regression at each silo. This general, iterative approach, known as Alternating Directions Method of Multipliers (11; 12), is guaranteed to converge (13), has suc-2 cessfully been applied to many problems (14; 15; 16), and for GWAS we show that moderately high accuracies can be achieved with a few iterations (also see (14)). We show that our pipeline is accurate, scalable, practical, and a significant improvement over the meta 1.0 approach.

Tangentially, our approach also confers an increased level of privacy to study participants recruited across multiple centers. This is an area of active research: recent concerns about data-7 sharing have lead to a growing body of literature and methods that offer distributed analysis. UNICORN (17) provides fast access to external controls matched to the population structure. Privacy-first GWAS methods usually fall into one of two camps: a) Homomorphic Encryption techniques like (18; 19) which let the analyst perform computation on encrypted data, or b) Differential Privacy techniques which let the analyst report results in a manner that obfuscates participant re-identification. Both families of methods are active areas of research but not the primary focus of this work.

## Results

We implemented an end-to-end canonical GWAS pipeline in our meta 2.0 framework and compared performance to both a mega-analysis and meta-study 1.0. We evaluated performance on datasets constructed from 2,270 European individuals in the POPRES dataset (20). Since POPRES does not have phenotype information, we simulate a case-control study by assigning random effect sizes to 10 SNPs according to a model similar to that of (21). Based on the overall risk, half of the individuals were assigned case status and the remaining half were assigned to the control group. See **??** for further details. Figure (2A) shows a Manhattan plot of the P values for the simulated data after correcting for the first 5 PCs. Figures 2B,C) show the projection of cases and controls onto the first 4 PCs. No geographical structure is observable that separates cases from controls (i.e. cases and controls are well matched).

**Figure 2:**
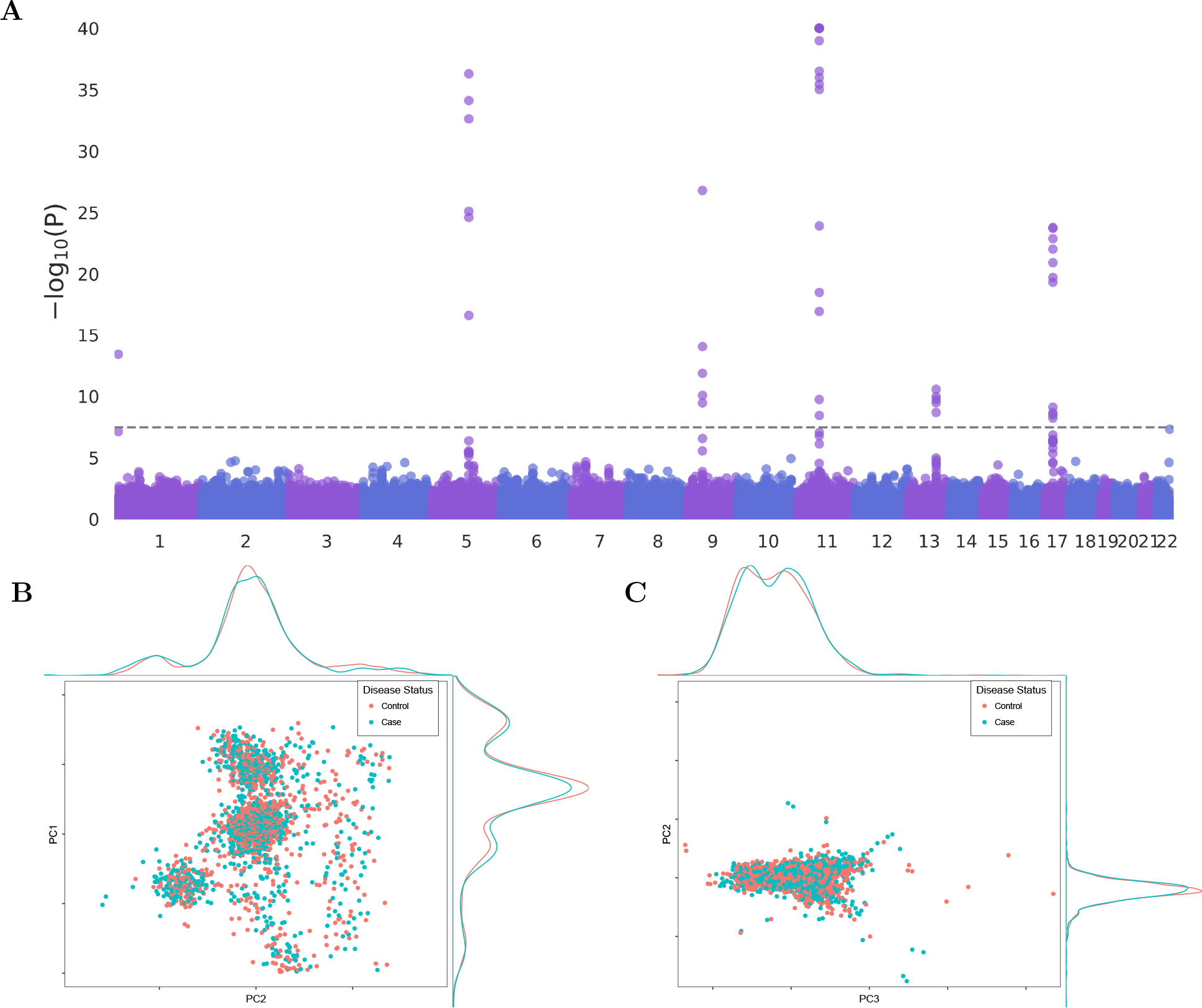
Simulated dataset. **A)** Manhattan plot of the simulated data. The P value are compute after centralizing the data and correcting for the first 5 PCs. The x-axis is SNP position and the y-axis is the significance on a − log_10_ scale. The dashed line marks the threshold for genome-wide significance (*P* = .01 after Bonferroni correction) (*n* = 2, 270). **B)** Projection of cases and controls onto the first two PCs. **C**) Projection of cases and controls onto the 3rd and 4th PCs

To evaluate our performance, we tackled two realistic scenarios. In scenario 1, we randomly split the data between 5 silos, simulating a fixed effect meta-study with a relatively small sample size per silo. In scenario 2, we split the data into only 2 silos based on case/control status, simulating a collaboration where cases and controls are recruited at different partner institutions. For each of these two scenarios, we compared performance using QC, PCA estimation, and PC-adjusted association analysis between our method and an equivalent centralized analysis. Lastly, to investigate the effects on extremely rare variants, we simulated a large, balanced, mega-cohort (*N* = 500, 000) and spread the data evenly among 10 data silos (collaborating centers).

### Step 1: Quality Control

The most important QC filters in a standard GWAS (call quality, missing-rate per individual, missing-rate per site, minor allele frequency, Hardy Weinberg equilibrium, and linkage disequilibrium) can be implemented efficiently under our paradigm. Call quality and missing-rate-per individual filters only require the communication of a threshold from the hub. Further, communicating (#missing, #heterozygous, #homozygous ref.) for each locus at each silo allows for computation of minor allele frequency (MAF), missing-per-loci and Hardy Weinberg equilibrium (HWE) at the hub. These values, along with the corresponding thresholds can be communicated back to perform MAF filters, missing-per-loci filters and HWE filters. Linkage disequilibrium (LD) pruning can also be implemented analogously, after communicating the local LD matrices from each silo to the center.

Table 1 demonstrates the comparison between our decentralized implementation and the centralized version of the equivalent filters as implemented by PLINK (22; 23). For each scenario, the table also includes the results of filtering based on local information at each silo (i.e. meta-analysis 1.0 approach). Furthermore, unlike the meta-study approaches, under our paradigm, relatedness of subjects across silos can be detected using locality sensitive hashing (24). Since our dataset does not contain related individuals and the performance of this approach has been previously analyzed by He, et. al. (24), we omit this filter.

**Table 1:**
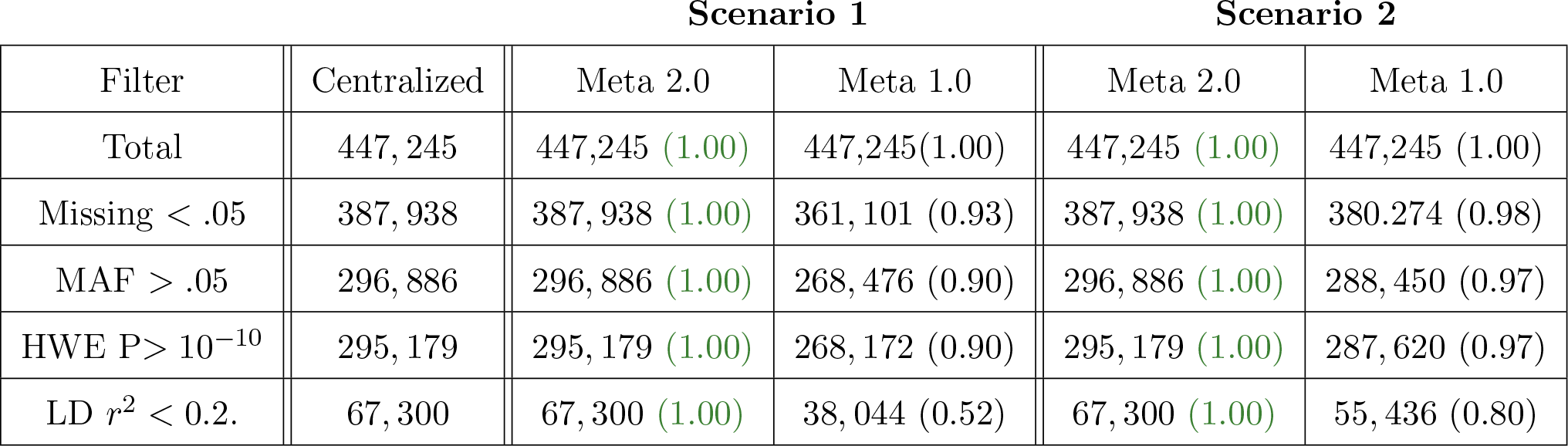
QC Comparison: The most important QC filters can be exactly implemented under a decentralized framework with no loss of accuracy. The table above contrasts our pipeline with both a centralized approach and a meta-analysis approach. For both scenarios under consideration results identical to the centralized approach can be achieved under the meta 2.0 framework. In this table, the number in parentheses is a measure of concordance (Jaccard index) between the set of unfiltered loci and the set of loci from the centralized pipeline. All centralized filters were implemented using PLINK(23; 22) (see **??** for details).

### Step 2: Population Structure Control

Correcting for the first few principal components (PC) has been shown to be an effective method of population structure control (25; 26). In this section we compare the results of a centralized and decentralized PCA on the POPRES dataset (20). First the data was filtered (MAF > 0.05, HWE with P value > 10^−10^, and LD of *r*^2^ < 0.2 using sliding windows of size 50 and strides of 25 loci). The remaining 67, 300 loci were normalized and algorithm 1 was used to perform PCA on the decentralized datasets. Figure 3 provides a visual comparison of the projection of samples onto the first two PCs in scenario 1. Projection onto higher PCs for both scenarios can be seen in Supplementary Figure **??** and Supplementary Figure **??** for scenario 1, 2 respectively.

**Figure 3:**
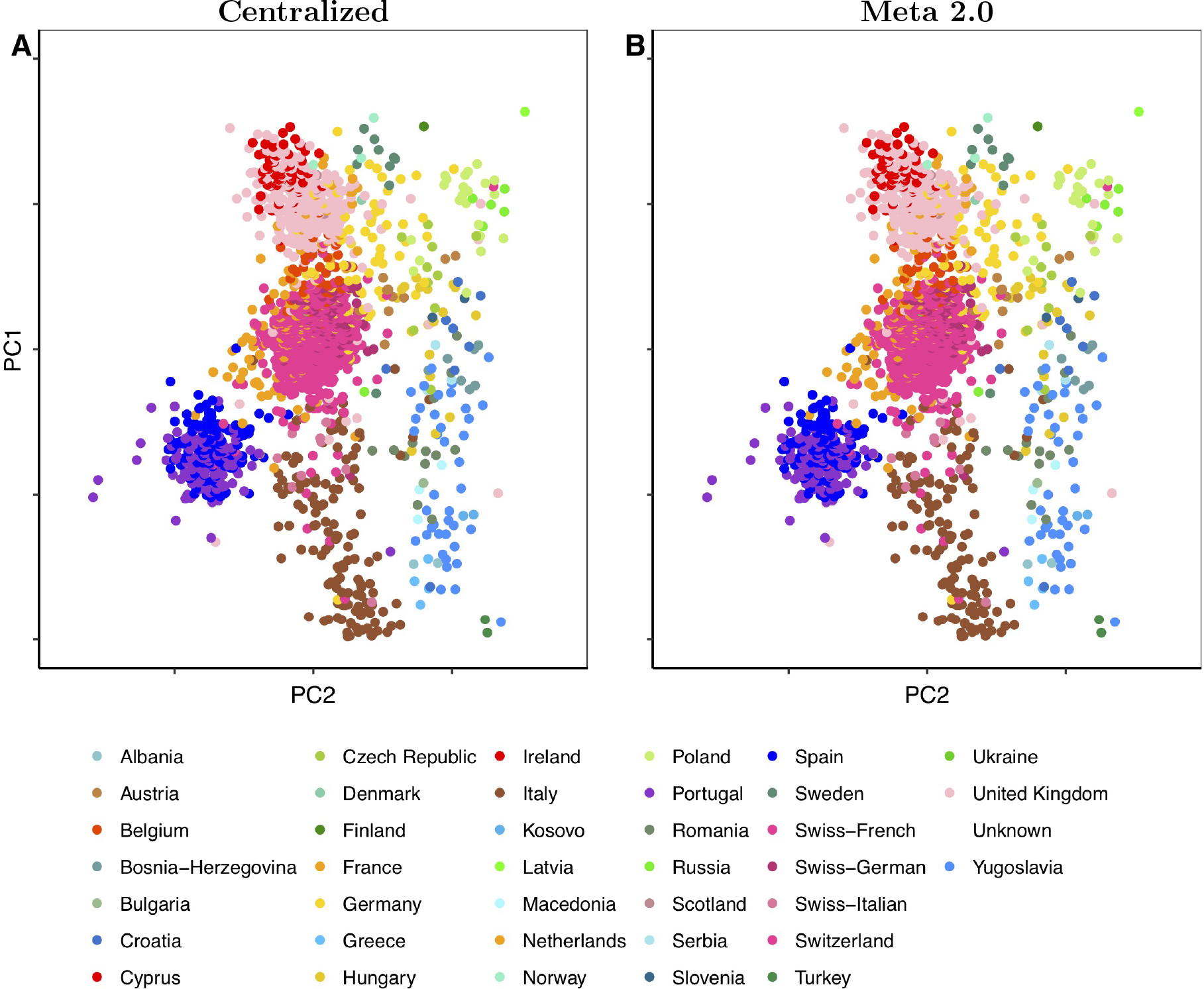
Principal components analysis comparison. Projection of POPRES individuals onto the first two principal components. **A)** Projection using singular value decomposition with centralized data **B)** Projection using singular value decomposition on decentralized data of scenario 1. The results are similar for scenario 2 and therefore have been omitted. See Supplementary Figure **??**, and Supplementary Figure **??** for projections onto higher PCs. Conventional meta-analyses cannot control for population structure using PCA of the global dataset

Communicating the LD matrices (SNP-SNP covariance matrix) constitutes the leading communication cost in algorithm 1. The size of this communication scales linearly in the number of silos and quadratically in the number of SNPs included and does not directly depend on the number of individuals in the study. In our dataset of 67, 300 post-QC SNPs, communicating the LD-matrix required a one-time 18GB transmission per silo (using 32 bit floats and without exploiting symmetry of the covariance matrix). The eigenvalue decomposition was performed using Implicitly Restarted Arnoldi Methods of ARPACK accessed through scipy (27; 28). On a single core machine, the decomposition for the top 5 vectors (using 15 Lanczos vectors) was performed in under 3 mins. While for the dataset under study, this is a significant time increase compared to local SVD decomposition, this algorithm does not scale with the number of individuals and presents a more competitive alternative for large-cohort GWAS studies.

### Step 3: Association

The final step of a typical GWAS pipeline is performing association analysis. This step often follows imputation of array data. Thanks to third-party imputation servers such as Michigan Imputation Server (29) and Sanger Imputation Server (30), fast, scalable, and reliable imputation is possible without centralizing the data. Under our decentralized paradigm, imputation would be performed by such a third-party service. Since the results are expected to be the same for centralized and decentralized data, in our pipeline we will omit this step and proceed to the association analysis.

We performed association analysis using logistic regression with the standardized value of each locus and the first 5 PCs as covariates. P values for each locus were computed using a likelihood ratio test (LRT) comparing the model containing only the first 5 PCs as covariates against the model that contains the SNP as well as the first 5 PCs. While the paradigm allows for computing P values using both Wald’s test and LRT, LRT was chosen as it tends to have higher power than Wald test (see Supplementary Figure **??**), particularly for rare variants in case-control studies (31). Additionally, LRT only requires each silo to communicate a likelihood value, where Wald test requires communication of the entire information matrix. Qualitatively, our results are similar for both tests (see **Supplement Figures** for all corresponding plots with Wald test).

In order to further probe our performance on very rare loci, we simulated an additional cohort with *N* = 500, 000. To simulate the cohort, an allele was created by randomly combining two haplotypes with 5 normally distributed nuisance random variables. These nuisance parameters account for 20% of overall variance in the burden and each loci accounts for 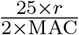 of the variance, where MAC is the minor allele count and *r* is a uniformly drawn random variable. Disease status was assigned to create a balanced study based on the overall burden.

Figure 2 **A**) shows a Manhattan plot for the P values of the POPRES cohort when the association is performed after centralizing the data. When the data is randomly distributed between the silos (scenario 1), a fixed-effect, meta-study approach can be taken to obtain parameter estimates and P values. Figure 4 **A** shows the concordance between parameter estimates of an inverse-variance fixed effect approach with those of a centralized study. Figure 4 **B** shows the corresponding plot for the P values. In this case, errors accumulate from, the difference in quality control, PC’s included in the regression and the uncertainty in the regression. In all cases, a cut off of 40 has been applied to − log_10_(P value). The meta-study approach appears to be slightly under-powered for highly significant loci. Figures 4 **C,D** compare the results of our fully decentralized pipeline where the regression parameters have been computed using an iterative approach. We observe a nearly perfect agreement between this decentralized approach and the centralized approach. The loss of power is further exaggerated when alleles under consideration have a low allele frequency. To probe this scenario we utilize our *N* = 500, 000 simulated dataset, described earlier. Figure 5 **A** compares the loss in power (Centralized logP - Decentralized logP) between an inverse variance meta-analysis and our approach (meta 2.0). While meta-analysis leads to a better initial estimate in all cases, within the first 5 iterations our approach out-performs these methods. Within 20 iterations, our approach effectively matches the power of the centralized analysis. As expected, when the minor allele count increases, both approaches perform better.

**Figure 4:**
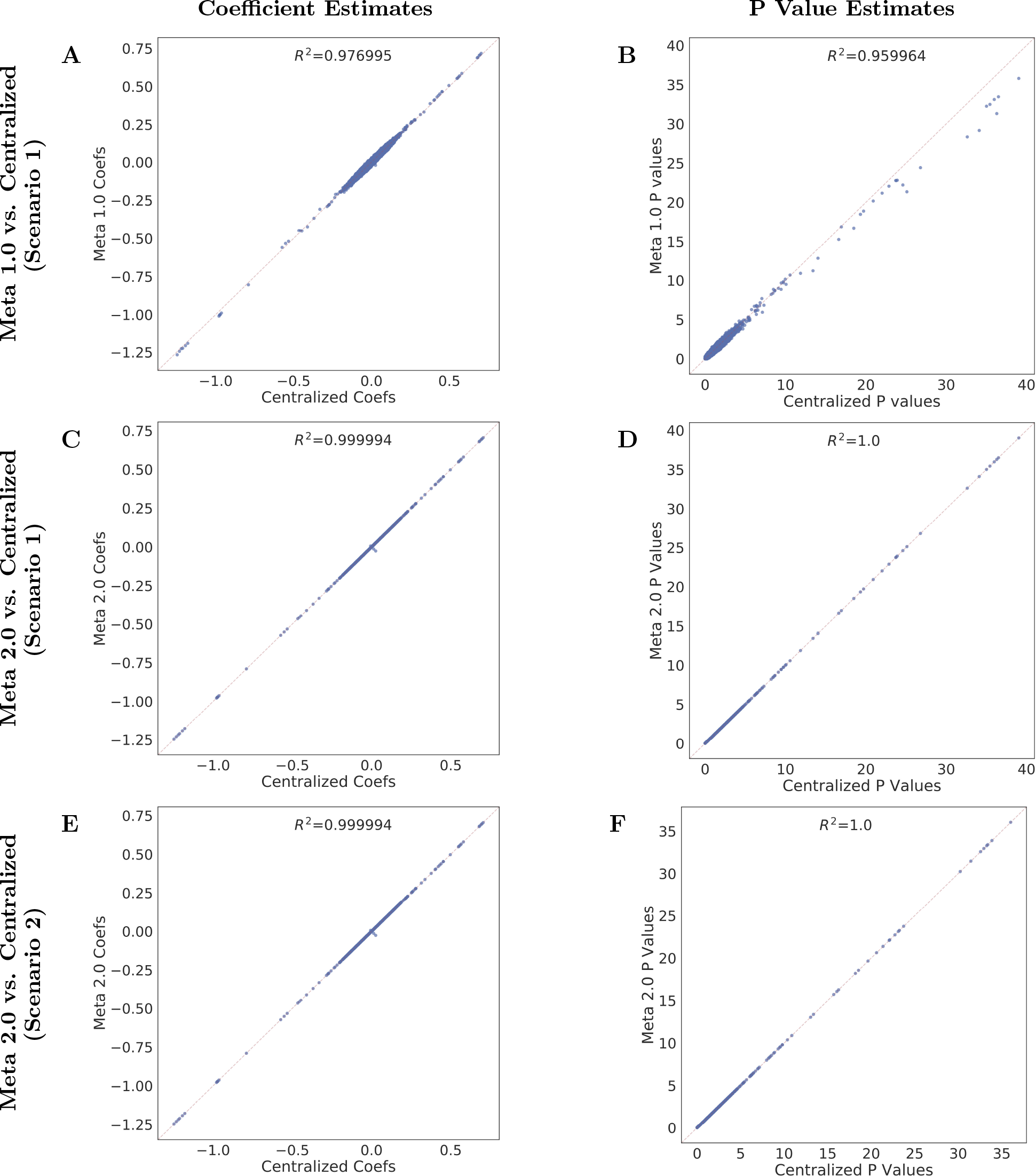
Association analysis. Concordance of effect sizes and − log_10_(P values) between centralized, meta 1.0, and meta 2.0 approach. In all figures a P value cut-off of 40 has been used. **A, B)** Concordance of a fixed effect approach with a centralized analysis in scenario 1. **C, D)** Concordance of our approach (100 ADMM iterations) with the centralized approach in scenario. **E,F)** Concordances between our method and a centralized approach in scenario 2 (100 iterations). In both scenarios our approach leads to results identical to a centralized analysis

**Figure 5:**
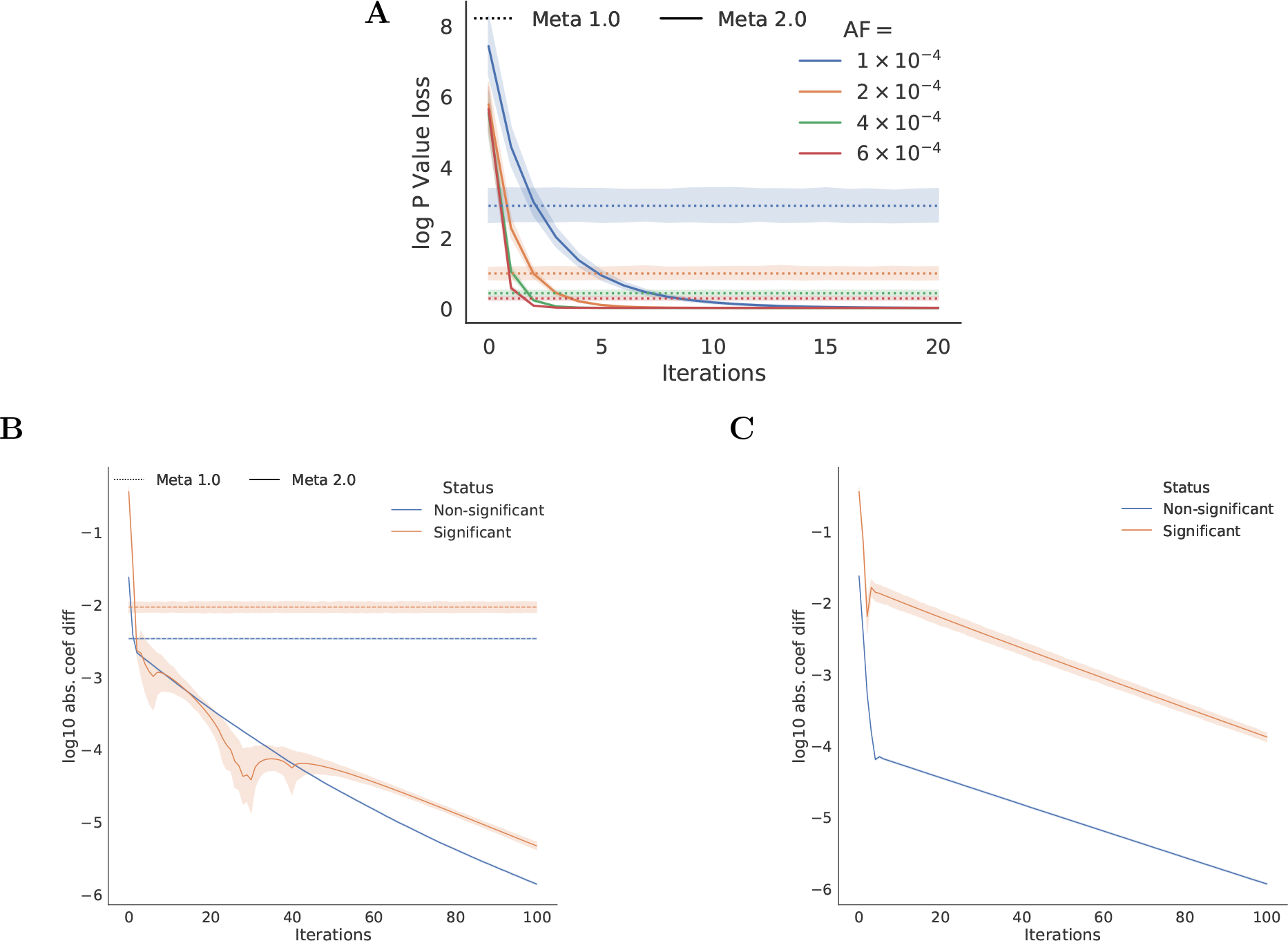
Convergence by iteration. For all plots, the x-axis shows the number of iterations of the algorithm (relevant only for our approach). **A**) Convergence of our approach for alleles with very low minor allele frequencies (MAF) in a large simulated dataset (*N* = 500, 000). The y-axis shows the difference in log10P value between a centralized study and one of the two decentralized approaches (100 draws). The minor allele frequencies correspond to minor allele counts of 50, 100, 200, 300. **B**) Convergence of the estimated coefficient to the centralized value for our approach and the inverse-variance FE approach, in scenario 1. **C**) Convergence of the estimated coefficient to the centralized value for our approach in scenario 2.

Scenario 2 challenges regression based meta-studies as each silo exclusively contains cases or controls, hence, local regression parameters cannot be computed. As figures 4 **E,F** show, our pipeline remains robust to this local label imbalance.

The decentralized regression is performed using alternating direction method of multipliers (32). Unless otherwise is specified, for each locus 100 iterations of the algorithm was performed. Each iteration requires the silo to communicate a local estimate of a regularized regression, and receive an updated estimate of parameter values (i.e. a back and forward communication of size equal to the number of covariates). Figure 5 **B, C** show the improvement in accuracy of the coefficient estimates (compared to the centralized estimates) as a function of number of iterations (data from chromosome 17, scenario 1 and 2 respectively). As before, in both cases good accuracy can be achieved in tens of iterations. The convergence for significant and non-significant loci is similar for scenario 1. In scenario 2, while the convergence rates are similar, significant loci lag behind non-significant loci. This is attributed to initializing the algorithm with 0, which is a better estimate for non-significant loci. Supplementary Figure **??** shows a similar analysis as a function of iterations and MAF. The centralized regressions were performed using sklearn’s LogisticRegression (*C* = 10^4^)(33). In our experiments, a serialized implementation of this approach (i.e. all local updates performed in serial and the global update performed after local updates from all silos), took approximately 3.7× and 1.5× longer than the centralized approach for scenario 1, 2 respectively.

## Discussion

The ability to perform full-fidelity association studies in a decentralized setting has logistical, security, and policy benefits when compared to centralizing the data. We believe this approach can lead to higher data security over the current paradigm where the data is often gathered by multiple institutions and subsequently centralized and reproduced at any institution approved for accessing the data. Our paradigm restricts the security and logistical risks to the few original data hosting centers. Securing these centers is cheaper, easier, and thereby more effective than securing every server where the accumulated data might be copied to. Furthermore, the reduction in the number of individuals with access to the raw data can reduce the privacy risks caused by human-error. Lastly, our algorithms only share aggregate level information, and unless the centers are case-only or control-only, they do not provide aggregate information by phenotype status. As a result, they act as a barrier against abuse and strictly improves on the privacy of sharing the raw data.

In contrast to meta-studies, which enjoy an increased level of the aforementioned benefits, our pipeline does not rely on large cohort asymptotic behavior at each silo, makes no assumptions regarding the distribution of phenotype status across silos, or uniformity of samples by population demography. While these parameters can effect the convergence rate of the association analysis, as demonstrated, under realistic scenarios (including complete separation of cases and controls) the convergence to relevant accuracies is fast. Furthermore, similar to centralized studies, our pipeline is flexible and can be extended to include new interaction effects, as well as other models (e.g. linear regression). The flexibility in choice of model, and weak requirements on the distribution of silos make our pipeline appealing for the future of GWAS.

These shortcomings in addition to concerns about data-sharing have lead to a growing body of literature and methods. UNICORN (17) provides fast access to external controls matched to the population structure - however it relies on knowledge about the population under study and is limited to control samples. Concerns about privacy of data sharing has brought about cryptographic GWAS approaches (18; 19). These methods allow for secure use of untrusted compute resources (e.g. cloud computation) but struggle for practicality and scalability. In particular, (18) is limited to simple computations and does not scale well. A more recent work (19) improves scalability and flexibility of analysis but still relies on a two step approach to limit the number of logistic regressions. These cryptographic approaches present a promising path forward but they are inefficient under a paradigm where the data is owned by each silo and the computations performed on the silos (and the hub) are not malicious. As a result, the homomorphic encryption literature is only tangentially relevant to our work. On the other hand, concerns about leakage of information from the reported results have given rise to differential privacy-preserving algorithms in genetics (34). With the expanding cohort sizes, the potential for leakage of individual-level information from the results decreases. The last broad area of relevant work is the decentralized/batch-based general machine learning algorithms that have become an ever-more-active area of research in order to meet the challenges presented by large datasets. Our work is inspired by, and can be expanded upon based on insights from these approaches (14; 35; 36; 37; 38; 39; 40; 41).

We see many opportunities to expand the current paradigm and pipeline. In this work we assume a minimal privacy threat model and focus on efficiency. One extension to our methods and paradigm would be to adopt an honest-but-curious threat model for all parties and aim to prevent leakage of individual level information from the data owning silos to any other party involved. Since for all steps of the pipeline the central hub simply sums the results of all silos (or takes averages), when more than 2 silos contribute, multi-party, secure sum protocols (42; 43) can be used to decrease the overall chance of information leakage. An alternative approach would be to replace the pipeline with differential privacy preserving algorithms. Indeed, many of the current components have differential privacy preserving counter parts (see (44; 45)). Yet another angle for expansion is to work in the regime where the number of silos and the cohort size are similar.

## Methods

Here we present the algorithms used to implement the fully decentralized GWAS pipeline.

### Notations

Unless otherwise is specified, we adopt the notations from table 2. Throughout, superscripts enumerate the silos.

**Table 2:**
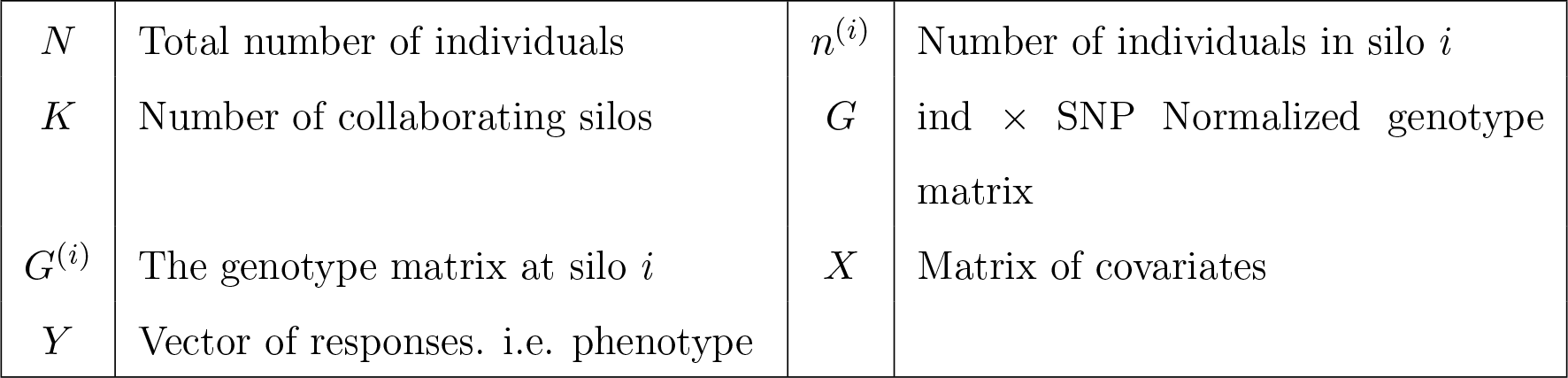
Table of symbols.

|  |  |  |  |
| --- | --- | --- | --- |
| $N$ | Total number of individuals | $n^{(i)}$ | Number of individuals in silo $i$ |
| $K$ | Number of collaborating silos | $G$ | ind × SNP Normalized genotype matrix |
| $G^{(i)}$ | The genotype matrix at silo $i$ | $X$ | Matrix of covariates |
| $Y$ | Vector of responses. i.e. phenotype | | |

### Data Filtering

A typical GWAS QC consists of many filters. The most important filters are easy to implement under the meta 2.0 framework. Missing-per-individual, quality-of-call filters and other individual level filters can be implemented as usual after a threshold is communicate from the central hub. After communicating the number of individuals per silo (*n*_*i*_), MAF, HWE, and missing-per-snp statistics for each loci can be computed using (# missing, # het., # homo. ref) per silo. Specifically, missing-per-snp and MAF statistics can be computed as follows:

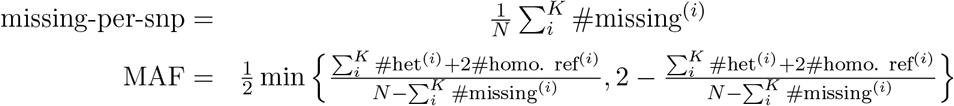

The HWE statistics is also a function of the total number of homozygous (ref and alt) and heterozygous individuals. For our experiments, we used the mid-p adjustment (46) as implemented by PLINK (22). After computing these statistics, filtering can be performed by simply communicating a threshold from the center to each silo.

The final filter we implemented is the LD-pruning filter. This filter is often used to choose sites for down-stream PCA analysis. In our implementation, we used a weighted averaging method similar to the one used for computing missing-per-snp statistics. However, whereas all the above filters require one back-and-forward communication of 𝒪(1) in size per locus, this filter requires a back-and-forward communication of order 𝒪(*w*^2^) per stride, where *w* is the LD-window size and an stride is the distance the pruning window moves per iteration. As a result, detection of LD moderately far away is difficult without a heuristic. Furthermore, whereas previous filters can be computed in chunks, and the computation on chunks can be parallelized, unless window size and stride size are the same, such optimizations are not possible under our current implementation. Nevertheless, these implementation inefficiencies are solely to reproduce the results provided by PLINK and can be avoided by real-world applicable heuristics.

Finally, we note that, as demonstrated by He et. al., it is possible to identify and filter for relatedness under the meta 2.0 paradigm.

### PCA

Ignoring population stratification can lead to false discoveries or a reduction in power(47; 48). Principal component analysis (PCA) is a common approach to reduce the adverse effects of stratification (47; 26). One common approach is to regress out the top PCs of the genotype matrix (26; 25). In this section, we discuss an implementation of such an approach.

Let *G* denote the normalized genotype matrix, with *G*_*i,j*_ represents the z-transformed frequency of the *i*^th^ individual at the *j*^th^ SNP. *G* = *U*Σ*V*^┬^ represents the singular value decomposition of *G*. Let *G*^(*i*)^ represent the corresponding matrix at silo *i* with all *G*^(*i*)^’s normalized using the global mean and standard deviation. Algorithm 1 describes the steps to perform a decentralized PCA. In this algorithm computing the full eigen-spectrum for *G*^┬^*G* can be prohibitively difficult. In real-world settings, PCA-based structure correction only relies on the top few eigen-components. Power iteration based methods, such as Lanczos algorithm, can be used to efficiently compute the top eigenvalue, eigenvector pairs. For this work we have used the implicitly Restarted Arnoldi Method (28) through the Scipy.sparse.linalg.eigsh interface (27).

#### Algorithm 1: Decentralized Principal Component Analysis

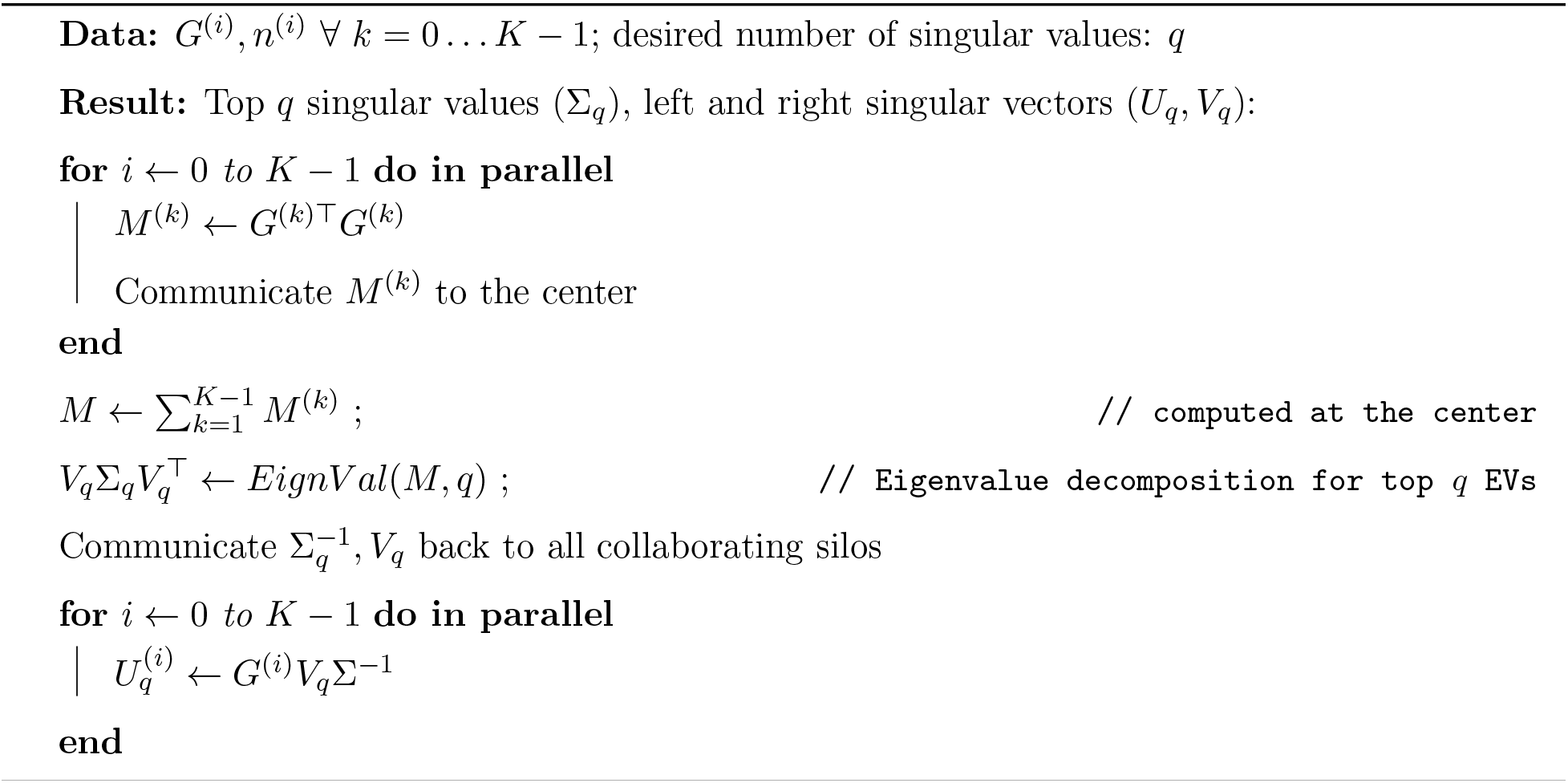

Algorithm 1 requires a one time communication of the gene-by-gene covariance matrix (*G*^(*i*)Τ^*G*^(*i*)^) from each silo to the center as well as a backward communication of 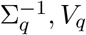. Typically we are interested in the top few PC’s, therefore, the backward communication to the silos is generally small; however, the covariance matrix can be relatively large (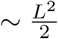 where *L* is the number of SNPs passing quality control for PCA). For 100k SNPs this would take an approximately 10GB communication per silo (using 32 bit floats and symmetry of the matrix). If the number of silos is very large, this communication can become overwhelming. In such a case, batch based approaches (41) or projection on the dimensions learned from a public dataset are viable alternatives.

### Association

Regression serves as one of the most common approaches in performing association in a GWAS. Under the maximum likelihood estimation (MLE) paradigm, performing a regression can be reduced to solving the following optimization problem:

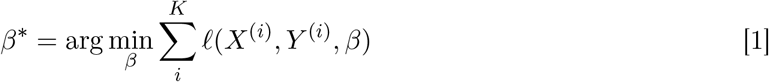

Where, *X*^(*i*)^, *Y* ^(*i*)^ are the matrix of covariates and vector of outcomes at silo *i* respectively, and 𝓁(*X*, *Y*, *β*) is the negative log-likelihood function. We use the alternating directions method of multipliers’ (ADMM)(14) framework to separate this problem by silos. To gain intuition, we explore the approach of inverse-variance fixed-effect meta-studies, where the MLE of each silo is provided (*z*^(*i*)^ = arg min_*x*_ 𝓁(*X*^(*i*)^, *Y* ^(*i*)^, *x*) and the meta-study estimate takes a weighted average of the local estimates:

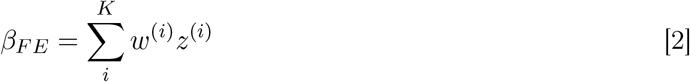

where *w*^(*i*)^ are a set of weights (inverse variance of the local estimate). With analogy to the fixed-80 effect approach the optimization problem of equation 1 can be written as:

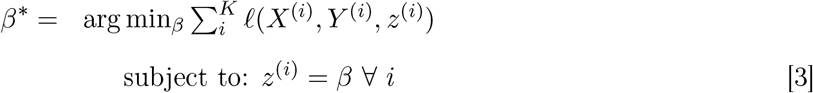

The constraint in equation 3 can be imposed using an augmented Lagrange(49; 50):

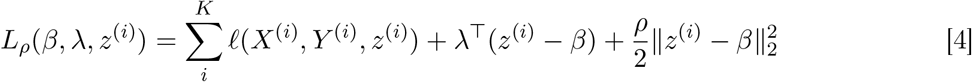

Here, λ is an estimate for the usual Lagrange multiplier and *ρ* is constant. This leads to the iterative solution

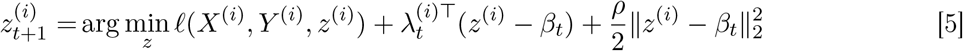

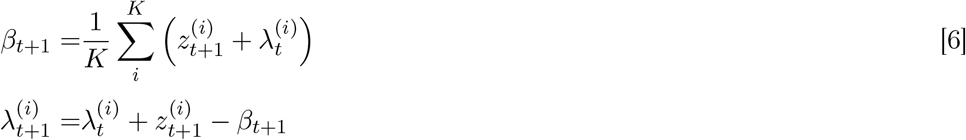

In words, at each iteration, all silos update their local estimates (*z*^(*i*)^) in parallel. These estimates are then communicated to the central hub where a consensus estimate is computed by averaging all local estimates. At the end of each iteration, this consensus parameter estimate is communicated back to each silo along with a silo specific dual variable λ^(*i*)^.

Note that in equation 6, after the first iteration, the update for the consensus estimate simplifies to 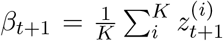. We contrast this equation to the fixed-effect, meta-study update of equation 2. Whereas fixed-effect updates use a weighted averaging that takes the variance of each local estimate (and therefore the sample size at each silo) into account, in our approach all the local estimates are simply averaged. The effects of sample size and the overall likelihood are taken into account when the local estimates are produced from the regularized regression of equation 5. For logistic regression, the likelihood function is given by 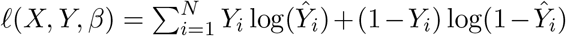 where *Ŷ*_*i*_ is the probability of a positive outcome for the *i*^th^ sample (i.e. 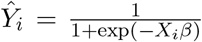. We used L-BFGS-B (51) with warm-start for the PCA estimates and default scipy(27) parameters to perform the update in equation 5.

### P-value computation

P values for the significance of each parameter can be constructed using both the likelihood ratio test (LRT) and Wald test. In this work we focus on LRT since it is both more efficient to compute and more powerful(31). To compute the P-value for a loci, each silo simply computes the local log-likelihood using the most recent global estimate. The local log-likelihood is then communicated and summed at the central hub to compute the overall log-likelihood. This global value can then be compared to the global log-likelihood of the model without the locus (which only needs to be computed once for the entire study).

To compute Wald test P values, the global parameter can be used to construct the local Fisher information matrix at each silo 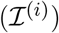. The overall information matrix can be assembled as the sum of these local matrices 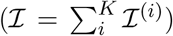. The Wald test P values can be computed using the overall information matrix at the central hub. Unfortunately, for a model with *p* parameters, this computation requires a communication of volume 𝒪(*p*^2^) per locus.

### Efficient implementation

For simplicity, we have presented all steps in the context of a single locus. In many cases, the network latency can be higher than a single computation. Fortunately, for most computations all loci are considered independently, therefore, it is possible to compute the updates on many loci and report them in groups. For instance, to compute the global allele frequencies, a vector of a few thousand local frequencies can be computed and reported at a time. Similarly, the ADMM updates on multiple loci can be computed and reported as a group. The silos can move on to compute the next block while awaiting the global updates.

## Code and Data Availability

POPRES data set is available through dbGaP. The preprocessing scripts and all other codes are available at https://github.com/apoursh/caringWithoutSharing.

## Acknowledgements

AP and CDB are supported by Genome Sequencing Project grant 1U01HG009080, Chan Zuckerberg Biohub. CDB and SP are supported by Clinical Genome Research Project grant U41HG009649. AP, CDB, SP are supported by CERSI grant U01FD004979.

